# *Escherichia coli* clonobiome: assessing the strains diversity in feces and urine by deep amplicon sequencing

**DOI:** 10.1101/735233

**Authors:** Sofiya G. Shevchenko, Matthew Radey, Veronika Tchesnokova, Dagmara Kisiela, Evgeni V. Sokurenko

## Abstract

While microbiome studies have focused on diversity on the species or higher level, bacterial species in microbiomes are represented by different, often multiple strains. These strains could be clonally and phenotypically very different, making assessment of strain content vital to a full understanding of microbiome function. This is especially important with respect to antibiotic resistant strains, the clonal spread of which may be dependent on competition between them and susceptible strains from the same species. The pandemic, multi-drug resistant, and highly pathogenic *E. coli* subclone ST131-*H*30 (H30) is of special interest, as it has already been found persisting in the gut and bladder of healthy people. In order to rapidly assess *E. coli* clonal diversity, we developed a novel method based on deep sequencing of two loci used for sequence typing, along with an algorithm for analysis of resulting data. Using this method, we assessed fecal and urinary samples from healthy women carrying *H*30, and were able to uncover considerable diversity, including strains with frequencies at <1% of the *E. coli* population. We also found that even in the absence of antibiotic use, *H*30 could complete dominate the gut and, especially, urine of healthy carriers. Our study offers a novel tool for assessing a species’ clonal diversity (clonobiome) within the microbiome, that could be useful in studying population structure and dynamics of multi-drug resistant and/or highly pathogenic strains in their natural environments.

**IMPORTANCE:** Bacterial species in the microbiome are often represented by multiple genetically and phenotypically different strains, making insight into subspecies diversity critical to a full understanding of the microbiome, especially with respect to opportunistic pathogens. However, methods allowing efficient high-throughput clonal typing are not currently available. This study combines a conventional *E. coli* typing method with deep amplicon sequencing to allow analysis of many samples concurrently. While our method was developed for *E. coli*, it may be adapted for other species, allowing for microbiome researchers to assess clonal strain diversity in natural samples. Since assessment of subspecies diversity is particularly important for understanding the spread of antibiotic resistance, we applied our method to study of a pandemic multidrug-resistant *E. coli* clone. The results we present suggest that this clone could be highly competitive in healthy carriers, and that the mechanisms of colonization by such clones need to be studied.

## INTRODUCTION

Microbiomes, both in terms of function and diversity, have recently been a topic of considerable interest. The gut microbiome has gotten special attention due to its high complexity and importance to health^1–9^. So far, studies have almost exclusively focused on species or higher-level diversity. However, this paints an incomplete picture, since strains within the same species can be of distinct clonal origin and have vastly different metabolic, pathogenic, and antibiotic resistance profiles^10–19^. Importantly, multidrug-resistant bacterial strains have been found competing with commensal strains in the gut, even without antibiotic pressure^18–23^. Thus, there is a pressing need to identify strains in the human microbiome for species of critical health importance.

*Escherichia coli* is one of the most common residents of the gut. While primarily a commensal colonizer, extra-intestinal pathogenic *E. coli* clones are implicated in a variety of diseases, including urinary tract infections (UTIs) - a leading cause of human antibiotic use^24–28^. The spread of multi-drug resistant *E. coli* is now a major health concern, especially the pandemic *fimH*30 subclone of sequence type ST131 (*H*30). Though recently-emerged, *H*30 is now globally distributed and comprises up to half of all urinary and bloodstream isolates of *E. coli* that are fluoroquinolone-resistant and produce extended-spectrum beta-lactamases (ESBL)^29–33^. Additionally, it is strongly associated with drug-bug mismatches and adverse outcomes in elderly and immunocompromised individuals^31–34^. Somewhat paradoxically, *H*30 is also a persistent gut colonizer of healthy people and frequently causes asymptomatic bacteriuria (ABU) in such carriers^35^. Yet, the relative clonal predominance of *H*30 strains among *E. coli* colonizing the gut or bladder in healthy carriers remains unknown. Answering these questions could have a significant impact on understanding the spread of antibiotic resistance and its reservoirs.

Currently, microbiome diversity is studied by sequencing the 16S rRNA gene, but this cannot capture clonal diversity^36, 37^. Conventional methods for assessing clonal diversity, such as metagenomic sequencing and single colony typing, are costly and labor intensive. For reliable clonal diversity analysis, metagenomic sequencing requires very high coverage per sample, while single colony typing requires handpicking large numbers of colonies for multi-locus sequence typing (MLST)^38–42^. In *E. coli*, MLST requires assessment of 7 genes per isolate which is analytically complex, costly, labor intensive, and therefore difficult to implement. Previously, we reported an alternative clonotyping method that requires sequencing regions of only 2 genes – *fumC* which is part of the MLST scheme and *fimH* that encodes a rapidly-evolving fimbrial adhesin^43^. The *fumC/fimH*-based (CH) typing of *E. coli* is widely accepted due to its simplicity and ability to not only identify specific STs but subdivide them into smaller subclones^43^. Specifically, *H*30 is identified using the allele combination *fumC*40*/fimH*30, while other less resistant ST131 strains have the same *fumC* but different *fimH* alleles.

Here, we report a high-throughput method for clonal typing of *E. coli* strains by combining CH typing and deep amplicon sequencing. We developed a new algorithm - Population-Level Allele Profiler (PLAP) - for detecting alleles and predicting the relative prevalence of each allele in a sample. We were able to assess the prevalence of clonal groups (including *H*30) in multiple fecal and urine samples concurrently, with a limit of relative abundance detection at <1% of the total population.

## RESULTS

### Deep amplicon sequencing of defined samples

To validate our approach and establish a limit of detection for strain presence, we first tested our deep amplicon sequencing procedure on a set of defined samples. To create the defined samples, we first selected a fecal sample from our lab collection known to contain *H*30 and ST101. Next, we isolated a single colony from each and confirmed them to be strains of *H*30 (*fumC*40/*fimH*30) and ST101 (*fumC*41/*fimH*86) using CH typing. From these single colonies, we first created *H*30-only and ST101-only mixtures of *fumC* and *fimH* amplicons. We also created four ST101/H30 mixed samples by combining the *fumC* and *fimH* amplicons from ST101 and H30 in ST101:*H*30 ratios of 1:1, 1:4, 1:100, and 1:1000.

Analysis of raw sequencing data from *H*30-only and ST101-only samples showed the average coverage of erroneous bases was 0.08% ± 0.09% for both strains. Erroneous bases were observed in both genes across most nucleotide positions. The highest coverage for an erroneous base was 0.66% of aligned reads in *fumC* and 0.45% in *fimH* for *H*30, and 0.68% in *fumC* and 0.46% of reads in *fimH* for ST101. The frequency distribution for erroneous base coverage is presented in Supplemental Figure 1.

Analysis of raw sequencing data from ST101/*H*30 mixes showed that both *H*30 and ST101 alleles were detectable in the 1:1, 1:4, and 1:100 mixes. In the 1:1000 mix, only alleles of the dominant *H*30 strain were observed. In the 1:1, 1:4, and 1:100 mixes, input and observed allele prevalence was highly correlated for both *fumC* and *fimH* (R^2^=0.996 and 0.997 respectively, Suppl. Fig. 2). Erroneous bases were observed at 0.09% ± 0.1% and 0.08% ± 0.09% of aligned reads in *fumC* and *fimH*, respectively (Suppl. Fig. 1). The highest coverage for erroneous bases among all mixes was 0.79% of aligned reads for *fumC* and 0.57% of aligned reads for *fimH*. Since 0.79% of aligned reads was the highest coverage for an erroneous base, we established 0.8% as a cutoff for correct base calling in both genes. This cutoff was used for all further PLAP analysis.

### Deep sequencing of study samples and allele prediction

Next, we applied PLAP to 67 participant samples (43 fecal and 24 urine) collected from a previous study^35^. A total of 128 *fumC* and 129 *fimH* alleles were predicted across all samples, of which 123 (96.1%) and 125 (96.9%) were previously known *fumC* and *fimH* alleles, respectively. 5 novel *fumC* and 4 novel *fimH* alleles were potentially detected. All novel *fumC* and *fimH* alleles were phylogenetically distant from other alleles predicted in the sample, indicating that these alleles are not artifacts of sequencing (Suppl. Fig. 3, 4). These novel alleles nonetheless clustered with other *E. coli fumC* and *fimH* alleles, indicating that these are novel *E. coli* alleles rather than alleles belonging to other species.

The average number of alleles predicted per sample was 1.91 ± 0.96 for *fumC* and 1.93 ± 1.01 for *fimH*. 43 samples had same numbers of predicted *fumC* and *fimH* alleles; 24 samples had different numbers of predicted *fumC* and *fimH* alleles (Fig. 1). Overall, the number of predicted *fumC* alleles correlated to the number of predicted *fimH* alleles with an R^2^ of 0.88 (Fig. 1).

**Figure 1.**
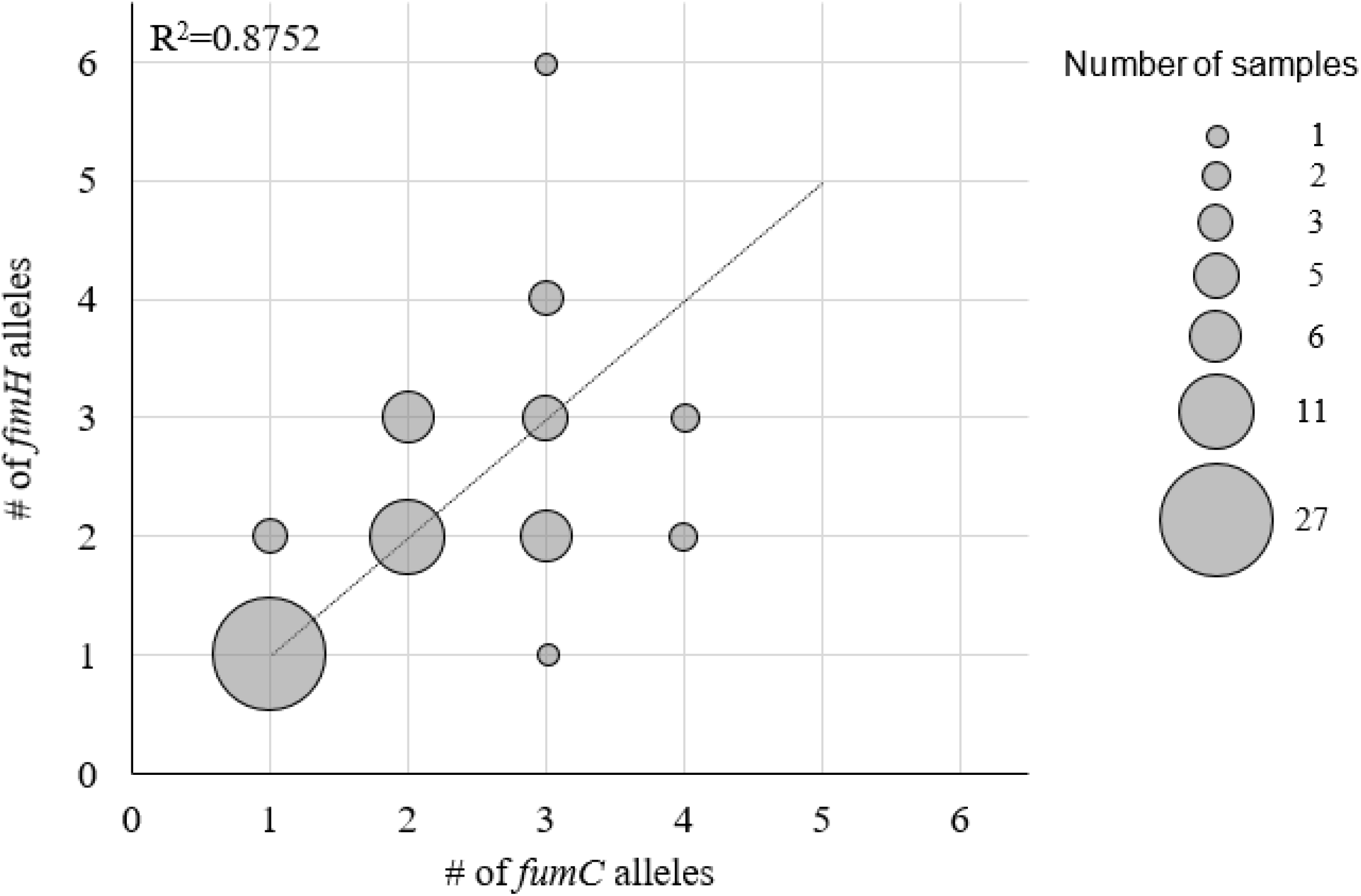
Congruency of *fumC* and *fimH* allele counts in fecal and urine samples. Size of bubbles corresponds to number of samples with designated *fumC*/*fimH* allele counts (i.e. 1 sample with one *fumC* allele and three *fimH* alleles). Linear fit with Pearson square correlation index shown.

To assess the performance of PLAP for predicting alleles, we used samples containing criterion clones - strains previously identified by single colony typing. PLAP detected criterion *fimH* and *fumC* alleles in 52 of these samples (90%). In the 6 samples where criterion allele(s) were not found, the criterion clones were ciprofloxacin-resistant, but their isolation from the sample required ≥2 plating attempts. This leads us to believe that these alleles were not detected because they were absent in the MacConkey-plated population prior to deep sequencing.

A total of 72 non-criterion (previously unidentified) *fumC* and 71 non-criterion *fimH* alleles were predicted by PLAP across all 67 samples. To assess the performance of PLAP on non-criterion alleles, we analyzed 14 samples (10 fecal, 4 urine) predicted to contain 22 non-criterion *fumC* and 22 non-criterion *fimH* alleles. 12 of these samples had at least one non-criterion allele alongside criterion alleles; the remaining 2 had multiple non-criterion alleles in each gene only. For each sample ≥40 single colonies were isolated and CH type determined using 7-SNP qPCR, with each CH type verified by sequencing. With these data, we confirmed 19 (86%) predicted non-criterion alleles for each gene. This included one predicted novel *fumC* allele. Of the unconfirmed alleles, one was not distinguishable by 7-SNP qPCR and had a predicted prevalence of 1%; therefore, we did not attempt to locate it. The remaining unconfirmed alleles had predicted prevalences of <3% and therefore may have been missed due to insufficient sampling. Additionally, all criterion alleles in these samples, 12 per gene, were predicted by PLAP.

### Prediction of allele prevalence in multi-allele samples

We have also designed PLAP to predict the within-sample prevalence of each allele. The average allele prevalence in fecal samples was 47.3% ± 4.3% SEM (range 0.88 – 100%) for *fumC* and 48.4% ± 4.22% SEM (range 1 – 100%) in *fimH*. The average allele prevalence in urine samples was 64.8% ± 6.91% SEM (range 1.4 – 100%) for *fumC* and 58.3% ± 7.18% SEM (range 1 – 100%) in *fimH*.

In order to verify that the prevalences predicted by PLAP were accurate, we compared predictions to actual in-sample prevalence using two different methods.

In the first method, we used *H*30 since ascertaining its prevalence is relatively simple. By plating the sample on MacConkey agar then patching onto LB-ciprofloxacin, it is possible to compare the number of cipro-resistant (*H*30) colonies to the total number of *E. coli* colonies. The ratio of these two numbers provides the *H*30 load in a sample. We compared the predicted prevalences of *fumC*40 and *fimH*30 to the *H*30 load in 17 fecal samples containing cipro-resistant *H*30. Correlations between the *H*30 load and the predicted prevalence of *fumC*40 and *fimH*30 were 0.86 and 0.84 respectively (Fig. 2), indicating that prevalences given by PLAP were representative of actual allele prevalences. To determine whether outliers were present, we calculated the 99% CI range for every sample (see Methods). Three outlier samples were identified (open circles, Fig. 2). Since it is possible that these outliers contain ciprofloxacin-sensitive non-*H*30 *fimH*30-containing clones, *fumC*-null or *fimH*-null clones, and/or ciprofloxacin-sensitive *H*30, we decided to employ screening of a large number of single colonies.

**Figure 2.**
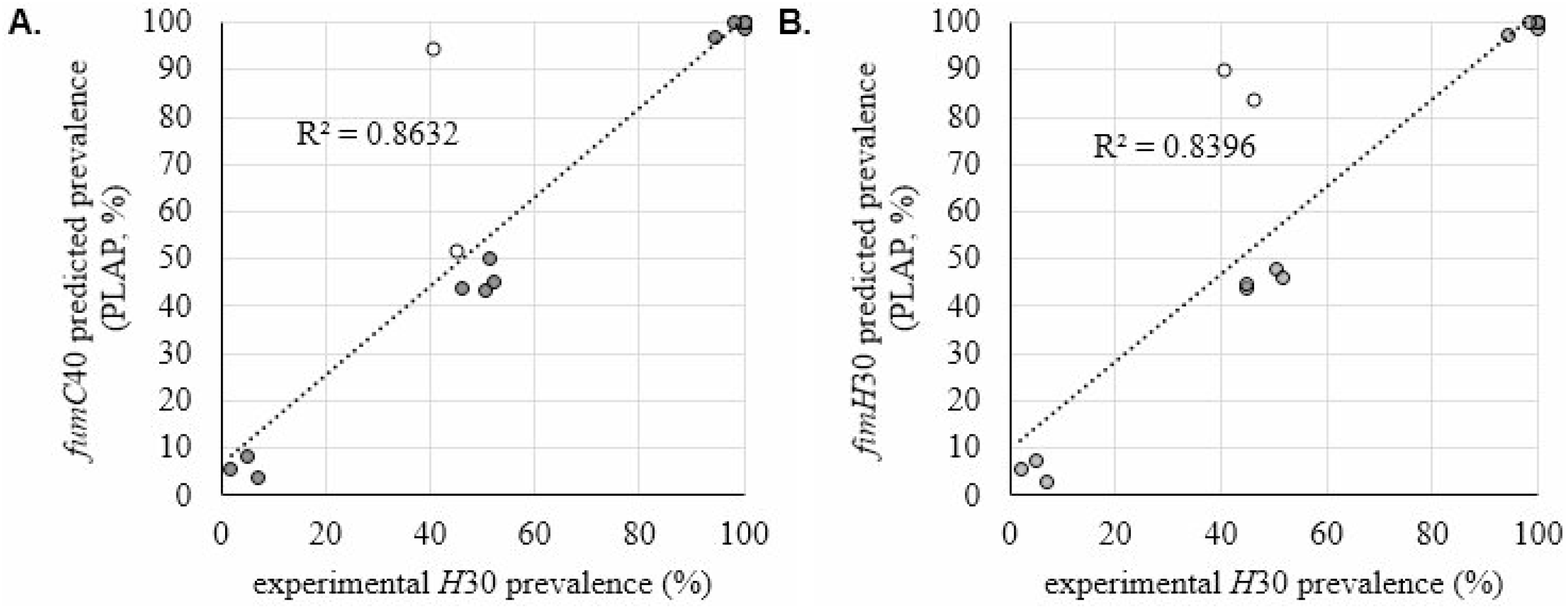
Validation of predicted *H*30 allele prevalence. PLAP-predicted prevalence of *H*30 alleles vs actual *H*30 load in *H*30-containing fecal samples. Prevalence of predicted *fumC*40 (**A**) and predicted *fimH*30 (**B**). Predicted prevalence of *fumC*40 and *fimH*30 is expressed as percentage of all *E. coli* in each sample. Experimentally confirmed *H*30 load is expressed as percent of *H*30 (ciprofloxacin-resistant) single colonies to all plated *E. coli* single colonies in percent. At least 130 colonies were tested per sample. Outliers, marked in open circles, were outside the 99% confidence interval of the number of colonies tested.

In this second method, we used single colony typing for the in-depth characterization of 14 multi-allele samples described above, alongside 4 additional single-allele samples (2 fecal, 2 urine) for which only one allele per gene was predicted. This set of 18 samples included 11 of the 17 fecal samples used for the *H*30-based analysis above, including one of the outlier samples. For all 18 samples, we used CH typing of ≥40 single colonies per sample to determine the prevalence of each *fumC* and *fimH* allele. Correlation between the PLAP-predicted prevalence and the experimental allele prevalence was 0.98 for both *fumC* and *fimH* alleles (Fig. 3). As in the *H*30 analysis above, we determined whether outliers were present using the 99% CI range for every sample. Only one outlier was detected, corresponding to the only sample that contained colonies from which *fimH* could not be amplified (*fimH*-null colonies). Furthermore, the sample that was an outlier in the *H*30-based analysis was found to contain a relatively rare ciprofloxacin-sensitive *H*30.

**Figure 3.**
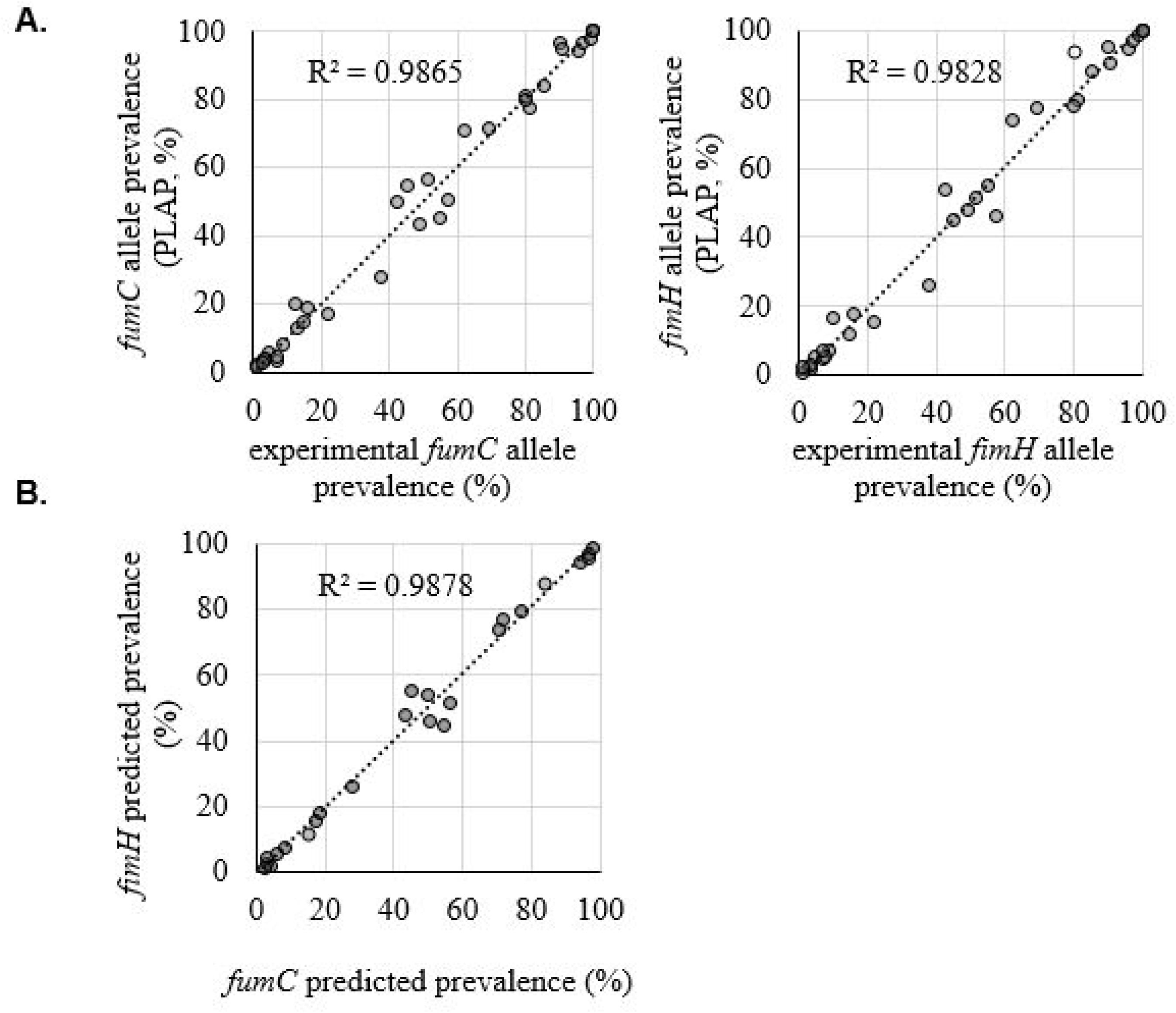
Validation of predicted *fumC/fimH* allele prevalence. **A.** PLAP-predicted vs experimental within-sample *fumC/fimH* allele prevalence in 18 samples. Experimental allele prevalence was determined by CH typing of at least 40 single bacterial colonies per sample. Outliers (open circles) were outside the 99% confidence interval of the number of colonies sampled. **B.** Predicted prevalence of *fumC* vs *fimH* alleles from the same CH type in 11 samples where no sharing of alleles between strains was present.

### Matching *fumC* and *fimH* alleles to predict sample strain content

In CH typing, unique combinations of *fumC* and *fimH* alleles are used to determine the identities of strains in a sample. Since a strain contains one copy of *fumC* and *fimH*, the prevalences of alleles of these two genes in the sequencing data should be identical. For example, in a sample containing 30% *H*30 (*fumC*40/*fimH*30) and 70% ST101 (*fumC*41*/fimH*86), we expect to see 30% of *fumC* reads to be *fumC*40 and 30% of *fimH* reads to be *fumH*30. In reality, however, the prevalences will be slightly different due to PCR and sequencing errors. To establish an acceptable difference between the prevalences of same-strain *fumC* and *fimH* alleles, we looked at 11 samples containing unique CH types (i.e. without allele sharing). In these 11 samples, the predicted prevalences of *fumC* and *fimH* were highly correlated (0.99, Fig. 3). First, we calculated the absolute difference between the predicted *fumC* and *fimH* prevalence for each matched pair of alleles. Next, each absolute difference was divided by the predicted *fumC* or *fimH* prevalence to obtain a relative deviation (Fig. 4). Finally, we used the relative deviations to derive an equation for the maximum acceptable difference between matching *fumC* and *fimH* alleles (Fig. 4).

**Figure 4.**
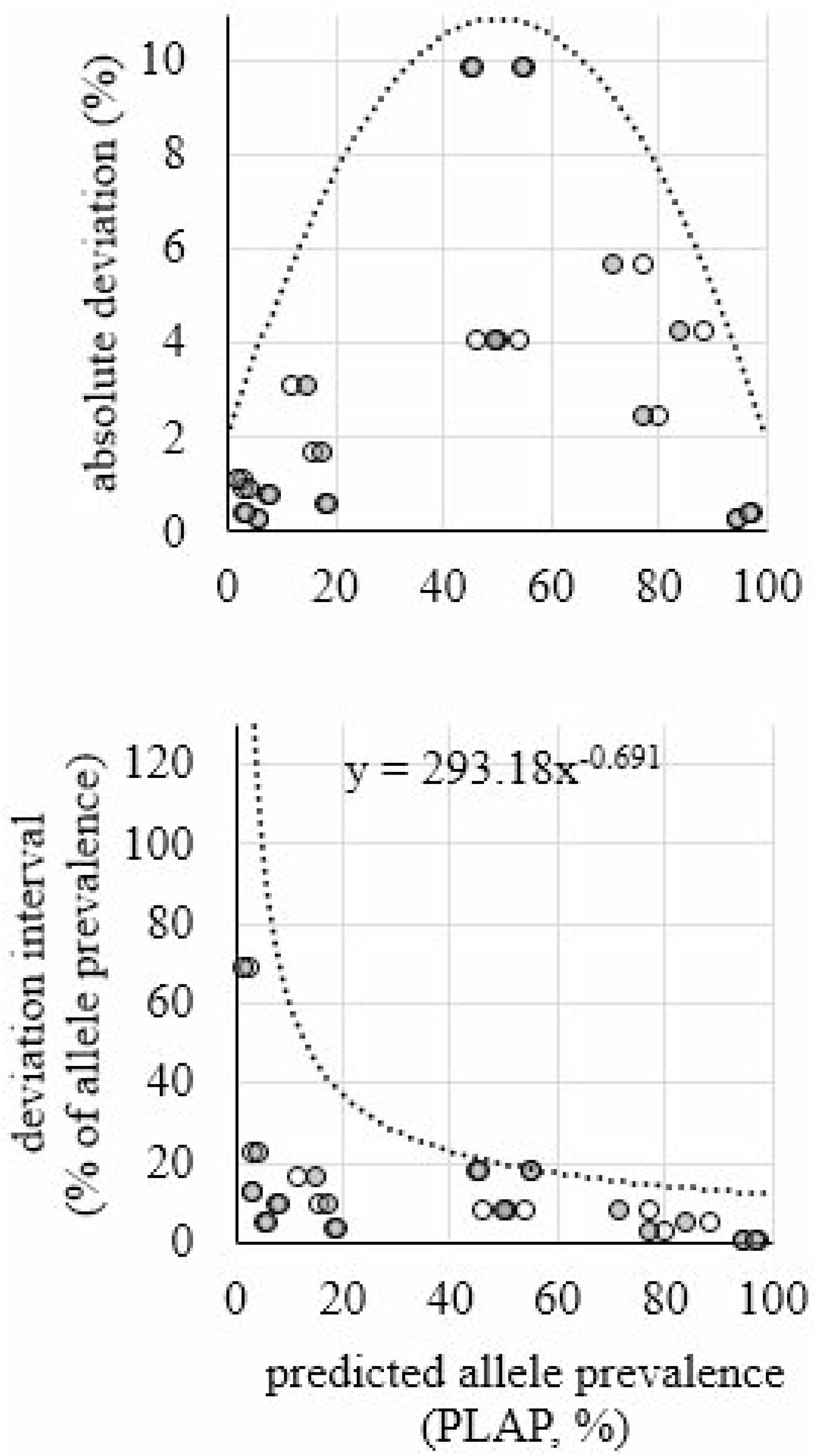
Difference in predicted prevalence between *fumC* and *fimH* alleles from the same *E. coli* strain. Deviation in absolute numbers is shown on the top. Deviation as a percentage of the prevalence of the allele is shown on the bottom. Open circles indicate *fimH* data points. Shaded circles indicate *fumC* data points. Trend lines and equations were used to determine intervals for matching (i.e. belonging to the same CH type) *fumC* and *fimH* alleles.

**Figure 5.**
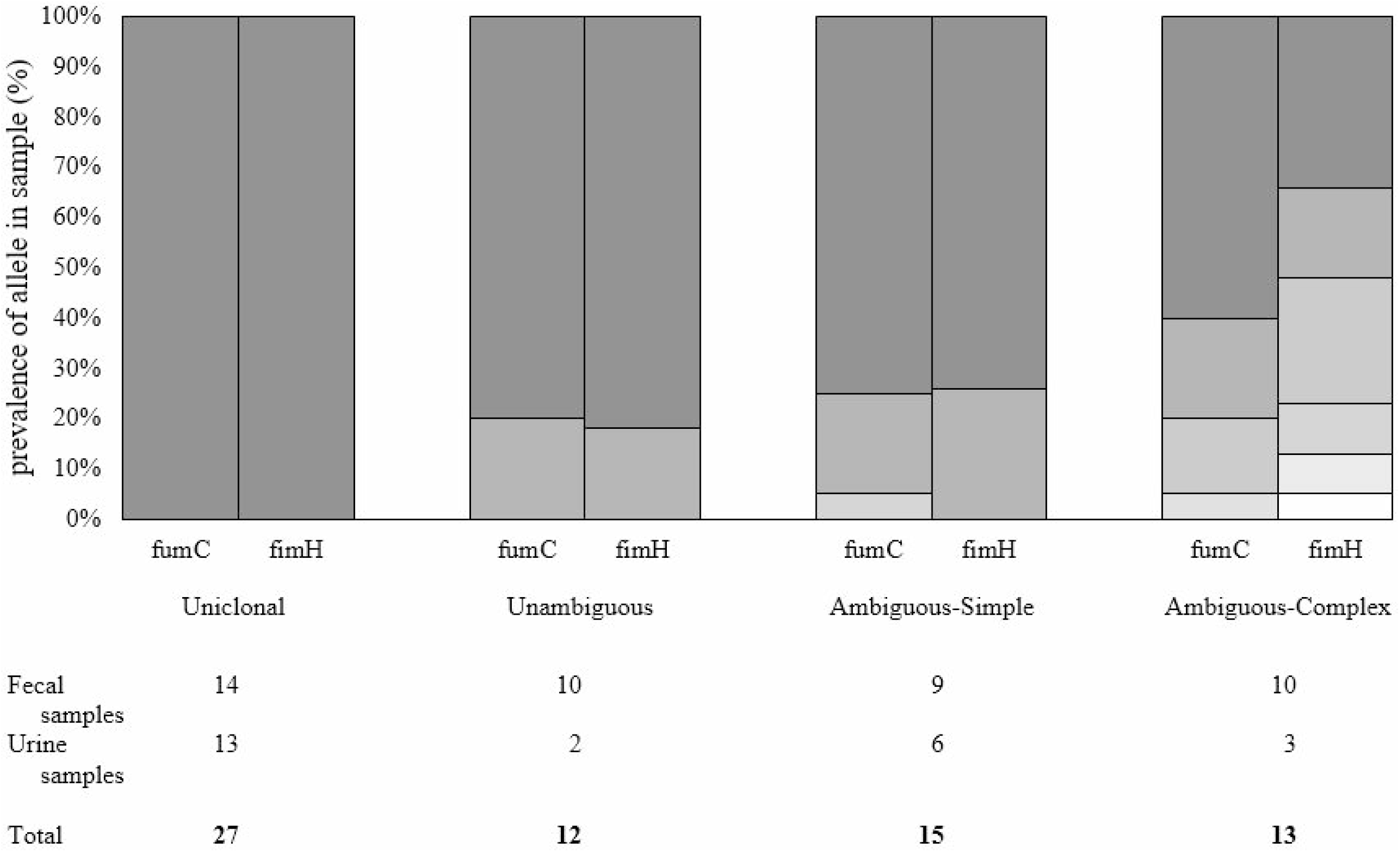
Representative examples of each sample category defined by within-sample breakdown of prevalence for *fumC* and *fimH* alleles. Number of fecal and urine samples belonging in each category is listed below.

While some samples, like those discussed above, contain only unique CH types, others contain CH types with shared alleles. For example, in a sample containing 30% *H*30 and 70% ST131, which share *fumC*40, the prevalence of *fumC*40 is not representative of either *H*30 or ST131 prevalence. For such samples, the minority rule was applied to resolve the strain content. Thus, under the minority rule, the percentage of *H*30 in the example above would be determined by *fimH*30, rather than *fumC*40, since the *fimH*30 prevalence is smaller. We tested this approach on both the *H*30 and the 18-sample analysis described above to see if this resolved outliers. In both cases, using the minority rule removed outliers and improved the correlation between predicted and experimental prevalence (Suppl. Fig. 5). Thus, we were able to assign strain content and strain prevalence in all samples, including samples with allele sharing.

### Predicted strain diversity of fecal and urine samples

Using the equation described above, we were able to classify all samples in our study into 4 categories (see Fig. 5): samples with only one CH type (uniclonal); samples with multiple unique CH types (unambiguous); samples with one dominant unique CH type and multiple minor non-unique CH types (ambiguous-simple), and samples where the dominant CH type was not unique (ambiguous-complex). Fecal samples were 33% uniclonal, 23% unambiguous, 21% ambiguous-simple, and 23% ambiguous-complex. Urine samples were 54% uniclonal, 8% unambiguous, 25% ambiguous-simple, and 12.5% ambiguous-complex.

Overall, 107 fecal and 48 urine strains were predicted, corresponding to 68 clones in fecal samples and 33 clones in urine samples. Of these clones, 50 (73.5%) and 24 (73%) were found in Enterobase, respectively.

Out of the 155 total strains predicted, 6 were *fumC*-null (3.9%) and 2 were *fimH*-null (1.3%). This is congruent with the occurrence of null alleles in our 18-sample subset, where 1 (3%) out of 35 total strains predicted was a null-allele strain.

The average number of strains per sample was 2.47 ± 1.32 for fecal samples and 1.96 ± 1.40 for urine samples. Based on Enterobase’s ST-phylogroup data, we determined that B2 was the most common (14 out of 47, 30%) among non-criterion fecal strains. Other phylogroups included A (26%), B1 (19%), C (8.5%), D (11%), E (2%), and F (4%). Non-criterion strains in urine samples included strains from phylogroups B2 (8 out of 16, 50%), B1 (19%), D (19%), A and F (6% each).

### Novel clones

17 fecal samples (40%) and 8 urine samples (33%) in our study were found to contain at least one novel CH type. This included 19 fecal and 9 urine CH types not found in Enterobase. Of these, 5 fecal and 3 urine CH types included at least one novel allele, and 14 fecal and 6 urine CH types were combinations of *fumC* and *fimH* that were not previously observed (novel CH combinations). Both CH types involving novel alleles and novel CH combinations were observed to be primarily low-frequency clones. The average predicted prevalence for novel CH combinations was 8.7% ± 3.5% SEM (range 1-64.2%), and 13 out of 20 novel CH combinations had predicted prevalences of <5%. One such combination was confirmed in our 14 characterized-sample set, consisting of *fumC*24 and *fimH*9, with a predicted prevalence of 1.6% and experimental prevalence of 1.2%.

Similarly, 7 out of 8 novel allele-containing CH types had predicted prevalences of <2%. The remaining CH type had a predicted prevalence of 70.7% and was detected using single colony typing. The novel *fumC* allele was paired with *fimH*47 and was verified to be 8 SNPs away from the closest known allele. The remaining MLST gene alleles for this strain were *adk*46, *icd*260, *mdh*160, *gyrB*266, *purA*1, and *recA*221.

### Clones below error threshold

To ascertain if we could identify alleles at prevalences below our defined error threshold of 0.8%, we ran PLAP on the set of 14 multi-allele samples using an error threshold of 0.5%. In 8 and 6 samples, respectively, prevalence of *fumC* and *fimH* alleles was <0.8%. None of the alleles corresponded to known *fumC* or *fimH* alleles. These apparent novel alleles clustered alongside known alleles identified in the sample (Suppl. Fig. 6, 7), leading us to conclude that these arose due to sequencing or amplification error rather than belonging to clonally different strains.

### Predicted strain diversity in urine and fecal samples

Strain diversity in first fecal samples was comparable with diversity in second fecal samples (paired t-test, p>0.1). Distinguishing between *H*30-containing and non-*H*30 samples showed that there was no statistical difference in strain diversity between *H*30-containing and non-*H*30 fecal samples of either kind (unpaired t-test, p>0.1), and that there was no difference in diversity between first and second fecal samples in either non-*H*30 or *H*30-containing samples (Fig. 6, paired t-test, p>0.1). Both *H*30 and non-*H*30 urine samples were less diverse than corresponding-fecal samples (paired t-test, p<0.01 and 0.02, respectively). However, *H*30 urine samples were less diverse than non-*H*30 urine samples (t-test, p=0.04).

**Figure 6.**
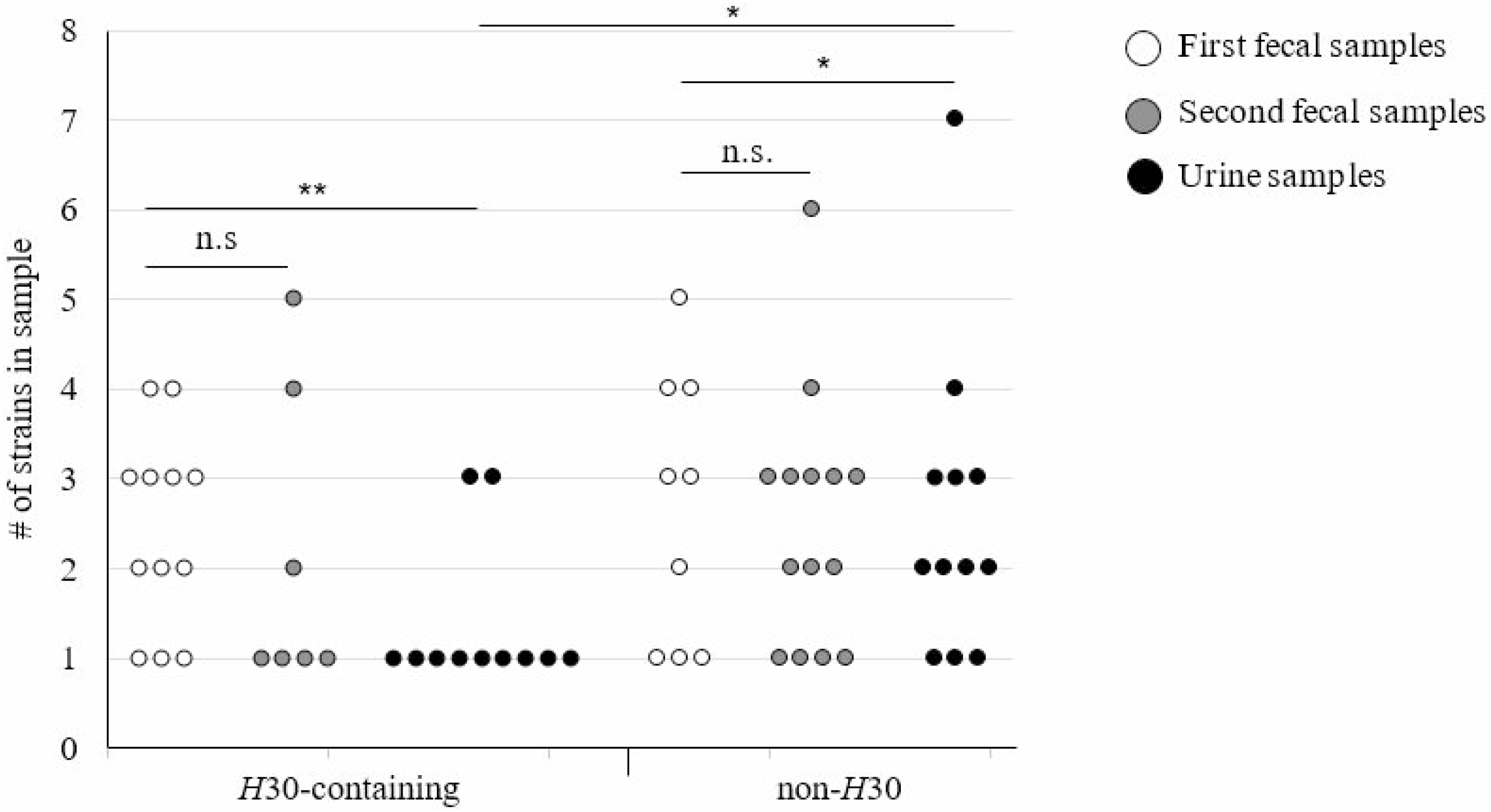
Diversity of *E. coli* in individual fecal/urine samples. *H*30 content was determined by PLAP and/or culturing.

It is also noteworthy that in 6 out of 23 *H*30-containing fecal samples, *H*30 was the only strain predicted, indicating that it may be fully dominant in the gut niche in these participants.

### Strain turnover in fecal samples

There was no correlation between number of strains in the first and second fecal sample, as well as no correlation between number of strains in the urine sample and either fecal sample (Fig. 7). When comparing the strain content of first and second fecal samples, we found that 92% of non-criterion strains appeared to be transient i.e. were detected in one of the fecal samples only. Transient non-criterion strains were also skewed towards lower-frequency strains (t-test, p<0.001, Fig. 8B). It is possible that these strains are present in both fecal samples but are below our limit of detection in one. However, we find that in one participant (P2, Suppl. Data) the first fecal sample contains 3 ciprofloxacin-sensitive non-criterion strains while the second fecal sample contains only ciprofloxacin-resistant *H*30 as verified by single colony testing. This leads us to believe that there may be significant strain turnover in our fecal samples overall.

**Figure 7.**
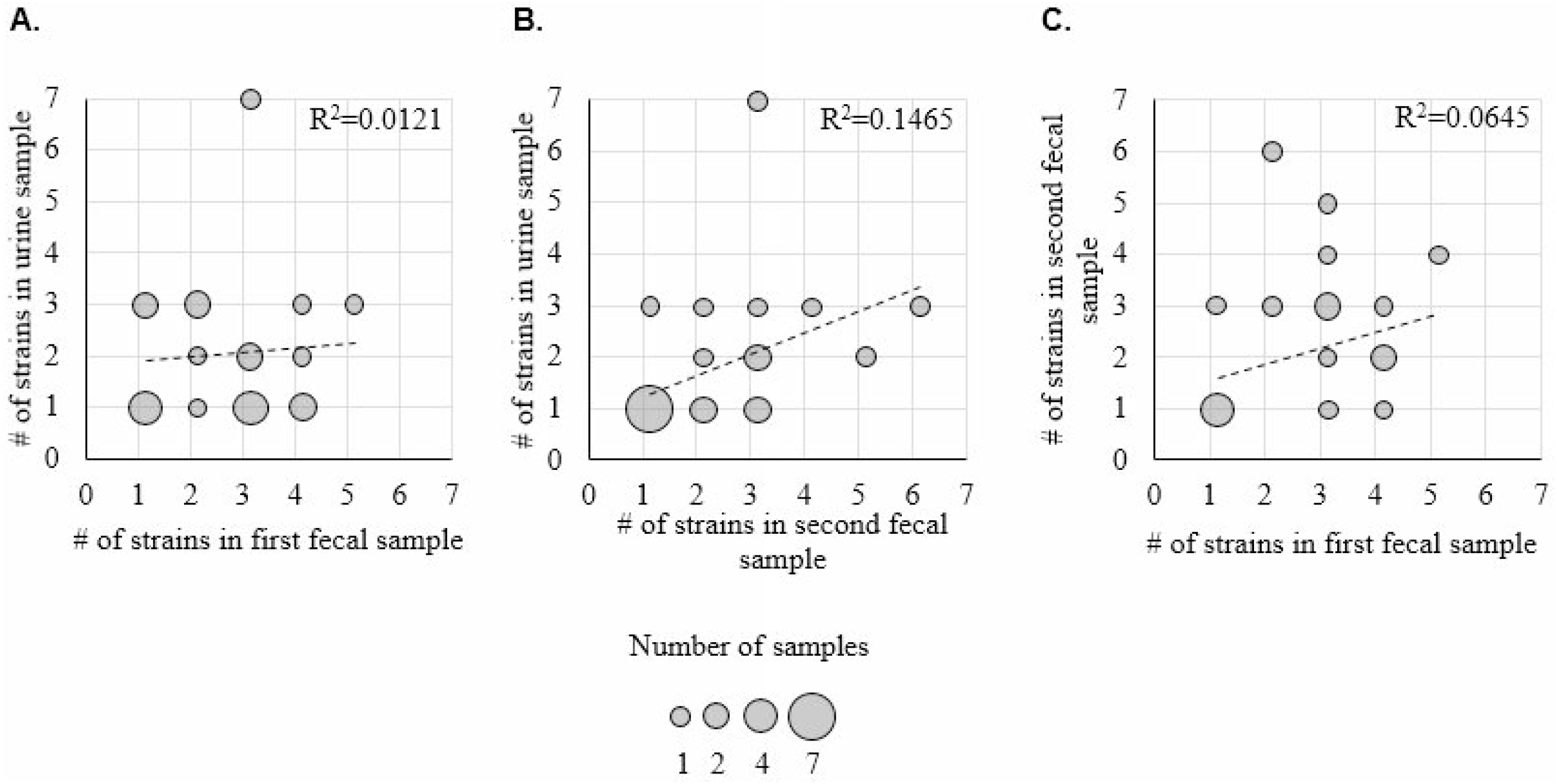
Counts of *E. coli* strains in fecal and urine samples. Number of strains detected by PLAP in (**A**) first fecal vs urine, (**B**) second fecal vs urine, and (**C**) first fecal vs second fecal samples. Each bubble indicates participants with the corresponding number of *E. coli* strains in the designated sample. The bubble size indicates number of participants with the determined number of strains. Linear fit with Pearson square correlation index shown.

**Figure 8.**
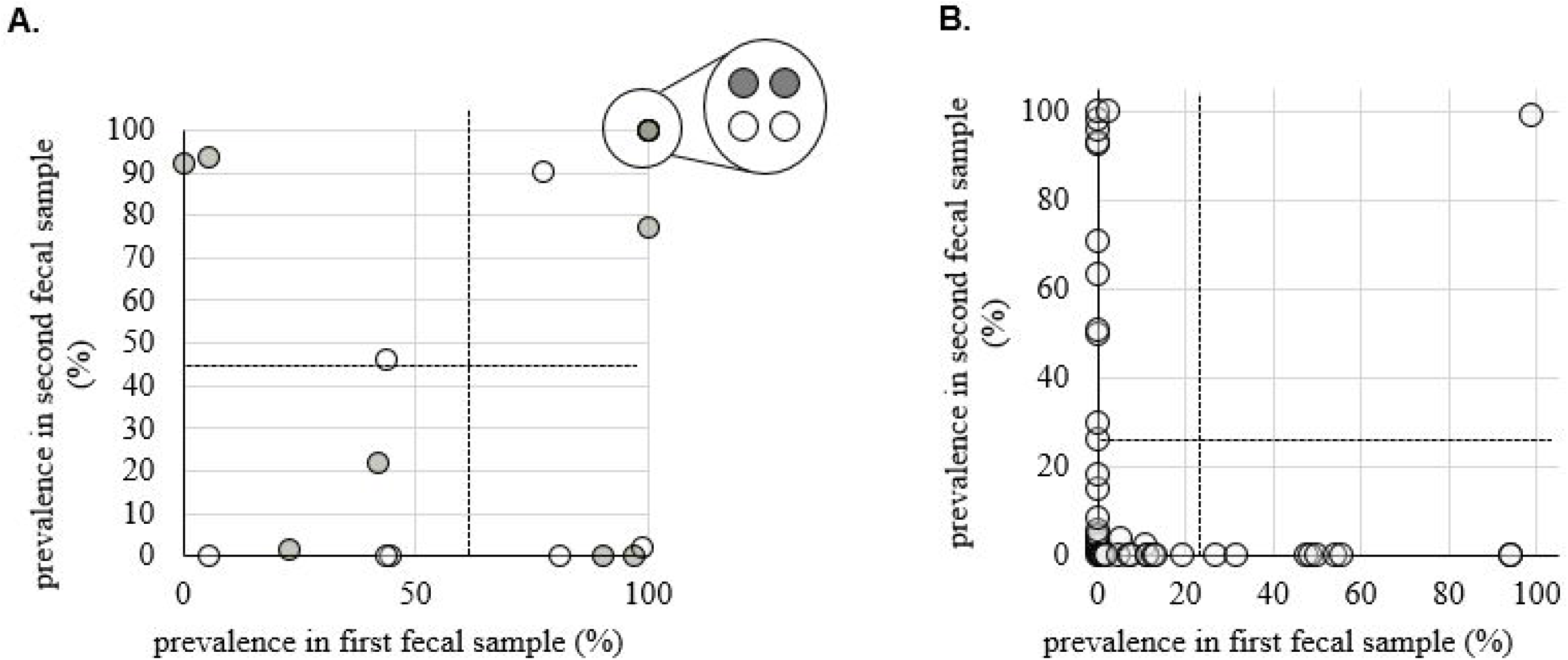
Persistence of *E. coli* strains in fecal samples. (**A**) Prevalence of criterion fecal strains in first vs second fecal samples. White data points indicate *H*30 strains while shaded data points indicate non-*H*30 strains. Circled cluster represents 4 strains present at 100% prevalence in both samples. Dotted lines indicate the mean prevalence for strains in first and second fecal samples. Distribution of prevalences in both first and second fecal samples is not significantly different from random (t-test, p>0.05). (**B**) Prevalence of non-criterion fecal strains in first vs second fecal samples. Dotted lines indicate the mean prevalence for transient strains in first and second fecal samples. Transient strains are defined as strains that are present in only one of the two fecal samples from the same participant. Distribution of prevalences in both first and second fecal samples is significantly skewed towards lower prevalences (t-test, p<0.01).

## DISCUSSION

We combined conventional *fumC/fimH* typing with deep amplicon sequencing to assess *E. coli* clonal diversity in a high-throughput manner. Our method has several advantages over existing protocols. Firstly, our method has high sequencing resolution for target species. Since we only sequence *E. coli fumC* and *fimH*, we can generate ≥0.5 million reads per sample, yielding ≥5,000 reads per base. In contrast, metagenomic sequencing, which is nonspecific to target species, yields only 20 reads per base per genome (assuming a 5Mb genome). Secondly, our method-assessed up to 46 samples per sequencing run. In contrast, MLST requires typing ≥100 single colonies per sample to capture the low-prevalence strains that PLAP detects. Finally, while we developed PLAP for *E. coli*’s CH typing, PLAP is not limited to *E. coli* clonotyping and may be generalized to other MLST schemes.

Despite studies showing that the healthy gut *E. coli* population typically includes multiple clones, we show that the pandemic multidrug-resistant subclone *H*30 can dominate the gut in healthy women, sometimes as the only detectable clone^42, 44–48^. This builds upon previous research which has found multidrug-resistant bacteria in healthy people, and healthy people who appear to harbor only one gut clone^44–48^. Total dominance is especially concerning since antibiotic pressure was absent, indicating that *H*30 is potentially outcompeting other clones by alternative means. Whether these mechanisms are metabolic, or whether certain virulence factors give *H*30 an advantage is unclear, though previous studies have speculated that some virulence factors may be beneficial for *E. coli* gut survival^49^. Additionally, our study involved a small number of participants in which *H*30 was present in the gut and bladder. Therefore, it is possible that host differences play a significant role. Another novel observation was that *H*30 was the sole detected urinary strain more frequently than other clones, regardless of *H*30 gut dominance/non-dominance. This may indicate that *H*30 might be an especially well-adapted uropathogen, potentially explaining its association with UTI. Since it is unknown how ABU converts to UTI, further study into *H*30 dominance in both ABU and UTI are needed.

We also uncovered substantial diversity in our samples. This includes significant *E. coli* diversity in non-*H*30 urine samples from healthy women. Reports of multi-strain bacteriuria are rare, likely due to the convention of selecting one isolate per urine sample^46, 47^. Therefore, it is unknown how common multi-strain bacteriuria may truly be. Remarkably, we also detected low-prevalence strains in the gut, some of which were novel clones, with up to 6 clones in a single sample. Gut *E. coli* diversity of this magnitude is supported by studies typing >200 single colonies per sample^42^. Studies using smaller counts usually report fewer clones, indicating that there may be undescribed *E. coli* diversity when manageable numbers of colonies are used^44, 45^. Therefore, we believe that microbiome-like approaches to *E. coli* diversity are necessary to fully understand intra-species dynamics in both the gut and bladder.

Our approach does have limitations. Firstly, our lowest detectable strain prevalence is 0.8% of the *E. coli* population. This limit may be addressed in several ways including use of a high-fidelity polymerase and preferential selection of *E. coli* colonies. However, we also recognize that detection of rare strains may still prove difficult and that methods like ours may not fully replace current techniques. Secondly, our method relies on sub-culturing *E. coli*. We are aware that, theoretically, some strains could be suppressed during growth on selective media, forming no/smaller colonies and skewing prevalence results. However, we did not encounter this during our study. While amplification of *fumC* and *fimH* may be applied to urine samples without culturing, attempts at doing this directly from fecal samples were unsuccessful, possibly due to *E. coli* comprising <1% of the gut microbiome, making *E. coli* DNA too rare to effectively amplify. Therefore, we used culturing for all samples. These issues lower the reliability of our approach, but we believe that it remains an important step towards development of comprehensive clonal diversity (clonobiome) assessment tools for any species of interest.

## MATERIALS AND METHODS

### Study design and sample processing

We selected a subset of participants from a previous study carried out by Kaiser Permanente Washington and University of Washington (Seattle, WA)^35^. That study identified healthy gut carriers of ciprofloxacin-resistant *E. coli*, including *E. coli H*30. These *E. coli* were found in initial fecal samples by plating on LB-ciprofloxacin and CH typing of 1 to 8 single colonies. After the initial fecal sample was analyzed, *H*30 carriers as well as carriers of some other strains were asked to provide urine samples. These were received on average 152 ± 55.9 days after the initial sample (85% responded). The respondents were then asked to provide follow-up fecal samples, which were received on average 82 ± 41.1 days after the urine sample (84% responded). All fecal and urine samples were tested for ciprofloxacin-resistant *E. coli* as with initial samples. For this study, we chose 28 individuals who supplied all three samples. In 11 participants, *H*30 was identified in all three samples; in 4 additional participants *H*30 was isolated in two samples. In 8 participants ciprofloxacin-resistant ST1193 was found in at least two samples. In 5 participants the same ciprofloxacin-susceptible clone was found in at least two samples. The sample types, strains clonal identity, and sampling times for all participants are shown in Supplemental Figure 8. Average age of participants was 66.7 ± 15.7 years.

### Preparation of predefined control samples

For control experiments, two predefined strains were chosen - *H*30 (*E. coli* FESS614.ds6) and clonal group ST101 (*E. coli* FESS614.ds4). DNA from these strains was extracted and *fumC* and *fimH* was amplified by PCR using the following conditions: 3min denaturation (95°C), 35 cycles of annealing (95°C for 45sec, 57°C for 45sec, 72°C for 45sec), 5min extension (72°C), 4°C hold. The primers (10 uM) used were as follows: 5’-TCACAGGTCGCCAGCGCTTC-3’ (*fumC* forward), 5’-GTACGCAGCGAAAAAGATTC3’ (*fumC* reverse), 5’-TCAGGGAACCATTCAGGCA-3’ (*fimH* forward), 5-ACAAAGGGCTAACGTGCAG-3’ (*fimH* reverse). Amount of PCR product was measured by Qbit. To create *H*30-only and ST101-only samples, the corresponding *fumC* and *fimH* PCR products were pooled together at a 1:1 ratio. To create mixes, *H*30 and ST101 amplicons of *fumC* were mixed together in ST101:*H*30 ratios of 1:1, 1:4, 1:10, 1:100, and 1:1000. The same was performed with *fimH* amplicons. The *fumC* and *fimH* mixes were then pooled together by ratio type to create mixes that had equal concentrations of total *fumC* and *fimH*. The DNA mixes were prepared for sequencing using Nextera XT DNA library prep kit using standard protocol. The resulting library was sequenced on the Illumina MiSeq (v3 kit). All mixes, except 1:10, reached coverage of ≥9,000X and were analyzed.

### Deep sequencing and allele analysis of the fecal and urine samples

Each fecal and urine sample was plated on MacConkey agar to reach ~1,000 *E. coli* single colonies per plate. All colonies were swabbed from the agar and DNA extracted using the Qiagen Blood & Tissue Kit. From this pooled DNA *fumC* and *fimH* genes were amplified by PCR by using the same primers and conditions as described above for control samples. Amplicons were then purified and pooled by sample using the Qiagen Gel Extraction kit, then prepared for sequencing using Nextera XT DNA library prep kit using standard protocol except for usage of 52.5ul of RSB in the final magnetic bead cleanup step. The resulting library was sequenced on the Illumina MiSeq (v3 kit). Sequencing data was analyzed using a Python program of our construction, Population-Level Allele Profiler (PLAP), and has been made available for public use on GitHub: github.com/marade/PLAP. The process is described below (see also Suppl. Fig. 9).

For each sample, adapter sequences were removed using Trim-Galore, and resulting trimmed reads were aligned to a list of all known *fumC* and *fimH* alleles using KMA with strict 99.99% identity matching^50, 51^. For each KMA-detected allele per sample, trimmed reads were again aligned to the sequence using Minimap2 and SAMtools^52, 53^. Any candidate allele which had at least 1 base supported by <0.8% of reads was removed from consideration. False positives were filtered using a moving 10bp window for each allele as follows. Reads of ≥100bp with 100% identity within the window were counted. Alleles with low initial coverage, unstable coverage (high average deviation from the mean), and high similarity in coverage pattern to an allele with more stable coverage were removed from consideration. If >3 alleles were left for consideration for a gene, 10bp moving window analysis was repeated with ≥200bp reads. If for any interval in this second analysis, >60% of coverage was lost compared to the first moving window coverage, the allele was discarded. Heterogeneity at any positions that remained undescribed by surviving alleles was recorded. Relative abundance of all alleles was determined using the minimum coverage found during first moving window analysis. In samples found by PLAP to be ≥50% made up of <100bp reads (overtagmented samples), allele prevalence was calculated manually by ascertaining base(s) unique to each allele and using the coverage of these base(s) to calculate prevalence.

Out of the 28 total sets of fecal and urine samples chosen for this study, at least one sample failed PCR amplification or sequencing library prep in 4 sets and therefore all samples from these sets were dropped. From the remaining 24 sets we were able to sequence *fumC* and *fimH* in all three samples. Out of those, 67 (89%) samples – 22 first fecal, 24 urine, and 21 second fecal – reached ≥9,000X coverage per gene and were included in the analysis.

### Determining within-sample clonal group breakdown

Identity of strains present in a sample was determined by combining *fumC* and *fimH* allele numbers and determining the ST type using Enterobase. In uniclonal and unambiguous samples, every allele had one match supported by the equation for maximum acceptable difference between same-strain *fumC* and *fimH*. Therefore, these alleles formed a CH type based on which ST type was determined.

For ambiguous-simple samples, the most prevalent *fumC* and *fimH* alleles formed an equation-supported CH type. Any alleles that also had a single equation-supported match were assigned to form a CH type. For all other alleles, Enterobase was consulted to determine which allele combinations have been observed. If the CH type(s) produced was between alleles that had different prevalences according to the equation, the “remaining” prevalence was calculated for the allele with the greater prevalence. This allele was then paired with allele(s) for which an Enterobase-logged CH type was not available and/or any novel alleles until the “remaining” prevalence was consumed. If there were any allele(s) that remained after this step, they were paired with the major allele of the opposite gene.

For ambiguous-complex samples, the most prevalent *fumC* and most prevalent *fimH* allele were assigned to the same CH type. The “remaining” prevalence was calculated for the allele with the greater prevalence and treated as an unmatched allele. From this step, we proceeded as with ambiguous-simple samples.

### Determining prevalence of clonal groups by culturing

Prevalence of ciprofloxacin-resistant clones in each sample was determined by diluting ~1ul of sample with ≥300ul of H_2_O, plating 40ul of this dilution on MacConkey agar, picking >130 single *E. coli* colonies, patching on Hardy-Chrom UTI agar to verify *E. coli* identity, then patching colonies on LB-ciprofloxacin. Prevalence of other clonal groups was validated by plating on MacConkey agar and subsequent patching of single colonies onto Hardy-Chrom UTI agar to distinguish *E. coli*. *fumC* and *fimH* alleles of these colonies were then determined by 7-SNP clonotyping and Sanger sequencing^54^.

### Statistical and phylogenetic analysis

To determine the 99% confidence interval (CI) for the prevalence of ciprofloxacin-resistant strains, the number of resistant colonies was treated as number of successes and the total number of picked colonies was treated as the total. To determine the 99% CI for the prevalence of ciprofloxacin-sensitive strains, the number of colonies of that strain was treated as number of successes and the total number of picked colonies was treated as the total. Confidence intervals were calculated using Stata^55^. All t-tests were run using GraphPad (http://www.graphpad.com/quickcalcs/ConfInterval1.cfm).

Phylogenetic trees were constructed using MEGA7^56^. Erroneous base coverage graph was generated using seaborn^57^. *Escherichia coli fumC* alleles were downloaded from Enterobase MLST allele data. *Escherichia coli fimH* alleles used are publicly available^58^. *Escherichia fergusonii* and *albertii fumC* alleles were downloaded from NCBI. *Klebsiella pneumonia* and *Enterobacter aerogenes* alleles of *fimH* were downloaded from the PATRIC database (www.patricbrc.org).

## ACKNOWLEDGEMENTS

We thank the personnel of KPWARI for assistance in collection of samples, and Dr. Sifang Chen for proofreading of the manuscript.

This work was supported by the National Institutes of Health (grant numbers R01AI106007 and R42 AI116114-02 [to E. V. S.])

E.V.S. conceived the project and designed the experiments. D.K. performed control sample sequencing and analysis. All other sequencing, validation, and analysis was performed by S.G.S. V.T. provided study data and samples. M.R. programmed the algorithm; M.R. and S.G.S. tested and calibrated it. S.G.S. and E.V.S. wrote the manuscript with input from all authors.

---

**Supplemental Figure 1**. Coverage of erroneous bases in *H*30-only, ST101-only, and mix sample sequencing. Coverage is expressed in percentage of total reads aligned to each gene.

**Supplemental Figure 2**. Correlation between input and PLAP-derived (deep seq) prevalences of *fumC* and *fimH* alleles of *H*30 and ST101 in 1:1, 1:4, and 1:100 mixes.

**Supplemental Figure 3**. Phylogenetic relationships between predicted novel *fumC* alleles and known *E. coli fumC* alleles. *Escherichia fergusonii* and *albertii fumC* alleles also presented for outgroup reference. Alleles not labelled with a species are known *E. coli* alleles or putative novel alleles. Alleles found in the sample as the novel allele are highlighted in the same color as the novel allele to show distance between predicted novel alleles and other *fumC* alleles present in the sample. Alleles present in multiple different samples are marked with the appropriate colors next to the allele name.

**Supplemental Figure 4**. Phylogenetic relationships between predicted novel *fimH* alleles and known *E. coli fimH* alleles. *Klebsiella pneumoniae* and *Enterobacter aerogenes fimH* alleles also presented for outgroup reference. Alleles not labelled with a species are known *E. coli* alleles or putative novel alleles. Alleles found in the sample as the novel allele are highlighted in the same color as the novel allele to show distance between predicted novel alleles and other *fimH* alleles present in the sample. Alleles present in multiple different samples are marked with the appropriate colors next to the allele name.

**Supplemental Figure 5**. **A**. Comparison of actual *H*30 load in *H*30-containing fecal samples to PLAP-predicted *fumC*-40/*fimH*-30 prevalences with minority rule correction (i.e. the smaller prevalence of the two was used). Prevalence of *fumC*-40/*fimH*-30 is expressed as percentage of all *E. coli* in each sample. *H*30 load is expressed as ratio of *H*30 (ciprofloxacin-resistant) single colonies to all plated *E. coli* single colonies in percent. **B**. PLAP-predicted allele prevalence (with minority rule correction) compared to experimental allele prevalence as determined by surveying at least 40 single colonies per sample.

**Supplemental Figure 6.** Putative rare novel *fumC* alleles identified by lowering the error threshold from 0.8% to 0.5%, marked in open shapes. Known alleles from the same sample as the rare novel allele are marked in filled-in shapes of the same type and color. FumC-40 was present in 3 different samples and therefore is marked by 3 different shapes.

**Supplemental Figure 7.** Putative rare novel *fimH* alleles identified by lowering the error threshold from 0.8% to 0.5%, marked in open shapes. Known alleles from the same sample as the rare novel allele are marked in filled-in shapes of the same type and color. FimH-30 was present in 3 different samples and therefore is marked by 3 different shapes.

**Supplemental Figure 8.** Sampling of volunteer sample sets. Length of segments is proportional to number of days between samples.

**Supplemental Figure 9. PLAP algorithm workflow.** Algorithms previously developed by other groups include Trim-Galore, KMA, Minimap2. Not pictured but used during windowed coverage checks is SAMtools.

